# Antifibrotic activity of a rho-kinase inhibitor restores outflow function and intraocular pressure homeostasis

**DOI:** 10.1101/2020.07.17.208207

**Authors:** Guorong Li, Chanyoung Lee, A. Thomas Read, Ke Wang, Iris Navarro, Jenny Cui, Katherine M. Young, Rahul Gorijavolu, Todd Sulchek, Casey C. Kopczynski, Sina Farsiu, John R. Samples, Pratap Challa, C. Ross Ethier, W. Daniel Stamer

## Abstract

Glucocorticoids are widely used as an ophthalmic medication. A common, sight-threatening adverse event of glucocorticoid usage is ocular hypertension, caused by dysfunction of the conventional outflow pathway. We report that netarsudil, a rho-kinase inhibitor, rapidly reversed glucocorticoid-induced ocular hypertension in patients whose intraocular pressures were uncontrolled by standard medications. Mechanistic studies in our established mouse model of glucocorticoid-induced ocular hypertension show that netarsudil both prevented and reversed intraocular pressure elevation. Further, netarsudil reversed characteristic steroid-induced pathologies as assessed by quantification of outflow function and tissue stiffness, and morphological and immunohistochemical indicators of tissue fibrosis. Thus, rho-kinase inhibitors act directly on conventional outflow cells to efficaciously prevent or reverse fibrotic disease processes in glucocorticoid-induced ocular hypertension. These data motivate a novel indication for these agents to prevent or treat ocular hypertension secondary to glucocorticoid administration, and demonstrate the antifibrotic effects of rho-kinase inhibitors in an immune-privileged environment.

## Introduction

Topical glucocorticoids (GCs) are routinely used after ocular surgery, and to treat common ocular disorders such as uveitis and macular edema (Noble and Goa 1998, Haeck, Rouwen et al. 2011). Unfortunately, ocular hypertension (OHT, i.e. elevated intraocular pressure [IOP]) is a common adverse event of such treatment, which can progress to a form of secondary glaucoma known as steroid-induced glaucoma. Interestingly, 90% of people with primary open-angle glaucoma (POAG), the most common type of glaucoma, develop OHT after GC treatment (Becker 1965). This is more than double the rate of the general population, likely because GCs increase extracellular matrix material, contractility and stiffness in an already dysfunctional conventional outflow pathway (Johnson, Gottanka et al. 1997, Zhou, Li et al. 1998), as is found in POAG (Ronkko, Rekonen et al. 2007, Tamm and Fuchshofer 2007). The conventional outflow pathway drains the majority of the aqueous humor, and its hydrodynamic resistance is the primary determinant and homeostatic regulator of IOP (Tamm 2009). Thus, pro-fibrotic changes in this outflow pathway, such as occur with glucocorticoid treatment, frequently lead to significantly elevated IOPs (Liu, McNally et al. 2018).

Rho-associated protein kinase (ROCK) is a major cytoskeletal regulator in health and disease. Indeed, ROCK mediates pro-fibrotic processes in many tissues and pathological conditions, including fibrotic kidney disease (Musso, De Michieli et al. 2017), idiopathic pulmonary fibrosis (Marinkovic, Liu et al. 2013, Knipe, Tager et al. 2015), cardiac fibrosis (Olson 2008, Haudek, Gupta et al. 2009)), liver fibrosis, intestinal fibrotic strictures associated with Crohn’s disease (Crespi, Dulbecco et al. 2020), vitreoretinal disease (Yamaguchi, Nakao et al. 2017) and lens capsule opacity (Korol, Taiyab et al. 2016). Hence, rho-kinase inhibitors (ROCKi) have been evaluated as anti-fibrotic therapeutics in multiple contexts. However, the mechanisms underlying their antifibrotic activity are complex and multi-factorial due to the central involvement of ROCK in many cellular processes. Thus, tissue/pathology-specific studies are essential to evaluate the efficacy of ROCKi’s as anti-fibrotic agents.

Mechanistic studies evaluating anti-fibrotic activity of ROCKi in the eye are particularly interesting in view of the eye’s immune-privileged status (Streilein 2003), which minimizes the role of macrophages and monocytes as compared to other organ systems. In fact, OHT in glaucoma patients is the only approved indication for ROCKi in humans thus far (Roskoski 2020). Specifically, two ROCKi, ripasudil and netarsudil (NT), have been approved for clinical use to treat OHT in glaucoma patients because of their safety profile and IOP-lowering ability and ultimately their unique mechanism of action (Garnock-Jones 2014, Isobe, Mizuno et al. 2014, Tanihara, Inoue et al. 2015, Kopczynski and Heah 2018, Serle, Katz et al. 2018). Therefore, ROCKi are the only available glaucoma drug class that directly targets, and improves conventional outflow function (Schehlein and Robin 2019). However, their use is currently limited to patients whose IOPs are inadequately controlled by medications that target ocular sites. Significantly, we here show that local delivery of the ROCKi, NT, dramatically lowers IOP in two cohorts of steroid-induced glaucoma patients refractory to conventional medications. Moreover, we define NT’s mechanism of restorative, antifibrotic action in treating steroid-induced OHT using our established mouse model (Li, Lee et al. 2019).

## Results

### NT lowered IOP in steroid-induced glaucoma patients whose ocular hypertension was poorly controlled by standard glaucoma medications

Based on changes to the trabecular meshwork (TM) previously observed in studies of steroid glaucoma (Johnson, Gottanka et al. 1997, Overby, Bertrand et al. 2014, Li, Lee et al. 2019), we tested NT’s efficacy at lowering IOP in steroid-induced ocular hypertensive patients who did not respond well to standard first line treatments. We retrospectively reviewed patient records, forming two cohorts of subjects from three treatment locations. The first cohort was created by an unbiased retrospective search of the Duke Eye Center’s electronic medical records using the key words, “steroid-responder/glaucoma” and “netarsudil”. Our search identified 21 eyes of 19 patients (mean age 66.8 years), treated with GCs for a variety of ocular conditions and who demonstrated ocular hypertension secondary to GC treatment (Table 1, Cohort 1). In accordance with current standard of treatment, these steroid-responsive patients were initially treated with aqueous humor suppressants (e.g., carbonic anhydrase inhibitors and/or adrenergics, Table 1). We note that prostaglandin analogues were typically used as second-line therapies (or not at all) in this patient cohort, due to concerns about the possible pro-inflammatory effects of these agents (Table 1). Unfortunately, these first and second line medications did not adequately control IOP in these patients, who presented with an average IOP of 24.3 ± 6.6 mmHg before NT treatment (mean ± SD). Thus, NT treatment was initiated, resulting in a clinically significant lowering of IOP in all patients within 1 month, presenting an average IOP decrease of 7.9 mmHg (p=1.2 × 10^−7^, Figure 1A). In fact, in this cohort, IOP was reduced to an average of 16.4 ± 4.9 mmHg, which is within the normal range. IOPs over 3 month course of treatment for each patient are shown in Figure S1. It is important to note that none of the patients were “tapered” from their steroid during the first month of treatment, and thus the observed IOP lowering cannot be due to a removal of steroid.

**Table 1:**
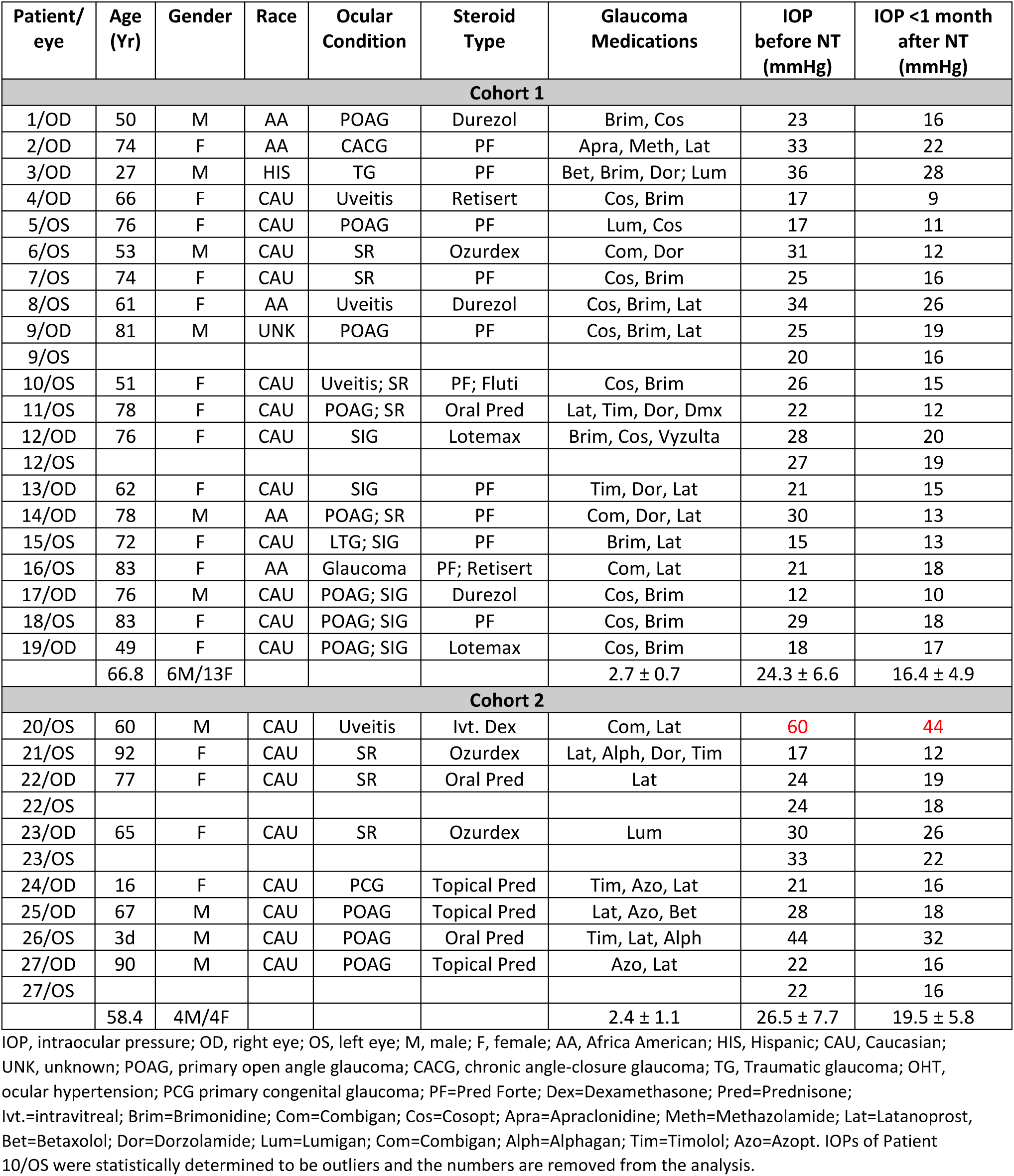
Netarsudil (NT) effectively lowered steroid-induced ocular hypertension in patients whose IOP was not well-controlled with standard glaucoma medications.

| Patient/<br>eye | Age<br>(Yr) | Gender | Race | Ocular<br>Condition | Steroid<br>Type | Glaucoma<br>Medications | IOP<br>before NT<br>(mmHg) | IOP <1 month<br>after NT<br>(mmHg) |
| --- | --- | --- | --- | --- | --- | --- | --- | --- |
| **Cohort 1** |  |  |  |  |  |  |  |  |
| 1/OD | 50 | M | AA | POAG | Durezol | Brim, Cos | 23 | 16 |
| 2/OD | 74 | F | AA | CACG | PF | Apra, Meth, Lat | 33 | 22 |
| 3/OD | 27 | M | HIS | TG | PF | Bet, Brim, Dor; Lum | 36 | 28 |
| 4/OD | 66 | F | CAU | Uveitis | Retisert | Cos, Brim | 17 | 9 |
| 5/OS | 76 | F | CAU | POAG | PF | Lum, Cos | 17 | 11 |
| 6/OS | 53 | M | CAU | SR | Ozurdex | Com, Dor | 31 | 12 |
| 7/OS | 74 | F | CAU | SR | PF | Cos, Brim | 25 | 16 |
| 8/OS | 61 | F | AA | Uveitis | Durezol | Cos, Brim, Lat | 34 | 26 |
| 9/OD | 81 | M | UNK | POAG | PF | Cos, Brim, Lat | 25 | 19 |
| 9/OS |  |  |  |  |  |  | 20 | 16 |
| 10/OS | 51 | F | CAU | Uveitis; SR | PF; Fluti | Cos, Brim | 26 | 15 |
| 11/OS | 78 | F | CAU | POAG; SR | Oral Pred | Lat, Tim, Dor, Dmx | 22 | 12 |
| 12/OD | 76 | F | CAU | SIG | Lotemax | Brim, Cos, Vyzulta | 28 | 20 |
| 12/OS |  |  |  |  |  |  | 27 | 19 |
| 13/OD | 62 | F | CAU | SIG | PF | Tim, Dor, Lat | 21 | 15 |
| 14/OD | 78 | M | AA | POAG; SR | PF | Com, Dor, Lat | 30 | 13 |
| 15/OS | 72 | F | CAU | LTG; SIG | PF | Brim, Lat | 15 | 13 |
| 16/OS | 83 | F | AA | Glaucoma | PF; Retisert | Com, Lat | 21 | 18 |
| 17/OD | 76 | M | CAU | POAG; SIG | Durezol | Cos, Brim | 12 | 10 |
| 18/OS | 83 | F | CAU | POAG; SIG | PF | Cos, Brim | 29 | 18 |
| 19/OD | 49 | F | CAU | POAG; SIG | Lotemax | Cos, Brim | 18 | 17 |
|  | 66.8 | 6M/13F |  |  |  | 2.7 ± 0.7 | 24.3 ± 6.6 | 16.4 ± 4.9 |
| **Cohort 2** |  |  |  |  |  |  |  |  |
| 20/OS | 60 | M | CAU | Uveitis | Ivt. Dex | Com, Lat | 60 | 44 |
| 21/OS | 92 | F | CAU | SR | Ozurdex | Lat, Alph, Dor, Tim | 17 | 12 |
| 22/OD | 77 | F | CAU | SR | Oral Pred | Lat | 24 | 19 |
| 22/OS |  |  |  |  |  |  | 24 | 18 |
| 23/OD | 65 | F | CAU | SR | Ozurdex | Lum | 30 | 26 |
| 23/OS |  |  |  |  |  |  | 33 | 22 |
| 24/OD | 16 | F | CAU | PCG | Topical Pred | Tim, Azo, Lat | 21 | 16 |
| 25/OD | 67 | M | CAU | POAG | Topical Pred | Lat, Azo, Bet | 28 | 18 |
| 26/OS | 3d | M | CAU | POAG | Oral Pred | Tim, Lat, Alph | 44 | 32 |
| 27/OD | 90 | M | CAU | POAG | Topical Pred | Azo, Lat | 22 | 16 |
| 27/OS |  |  |  |  |  |  | 22 | 16 |
|  | 58.4 | 4M/4F |  |  |  | 2.4 ± 1.1 | 26.5 ± 7.7 | 19.5 ± 5.8 |

IOP, intraocular pressure; OD, right eye; OS, left eye; M, male; F, female; AA, Africa American; HIS, Hispanic; CAU, Caucasian; UNK, unknown; POAG, primary open angle glaucoma; CACG, chronic angle-closure glaucoma; TG, Traumatic glaucoma; OHT, ocular hypertension; PCG primary congenital glaucoma; PF=Pred Forte; Dex=Dexamethasone; Pred=Prednisone; Ivt.=intravitreal; Brim=Brimonidine; Com=Combigan; Cos=Cosopt; Apra=Apraclonidine; Meth=Methazolamide; Lat=Latanoprost, Bet=Betaxolol; Dor=Dorzolamide; Lum=Lumigan; Com=Combigan; Alph=Alphagan; Tim=Timolol; Azo=Azoft. IOPs of Patient 10/OS were statistically determined to be outliers and the numbers are removed from the analysis.

**Figure 1:**
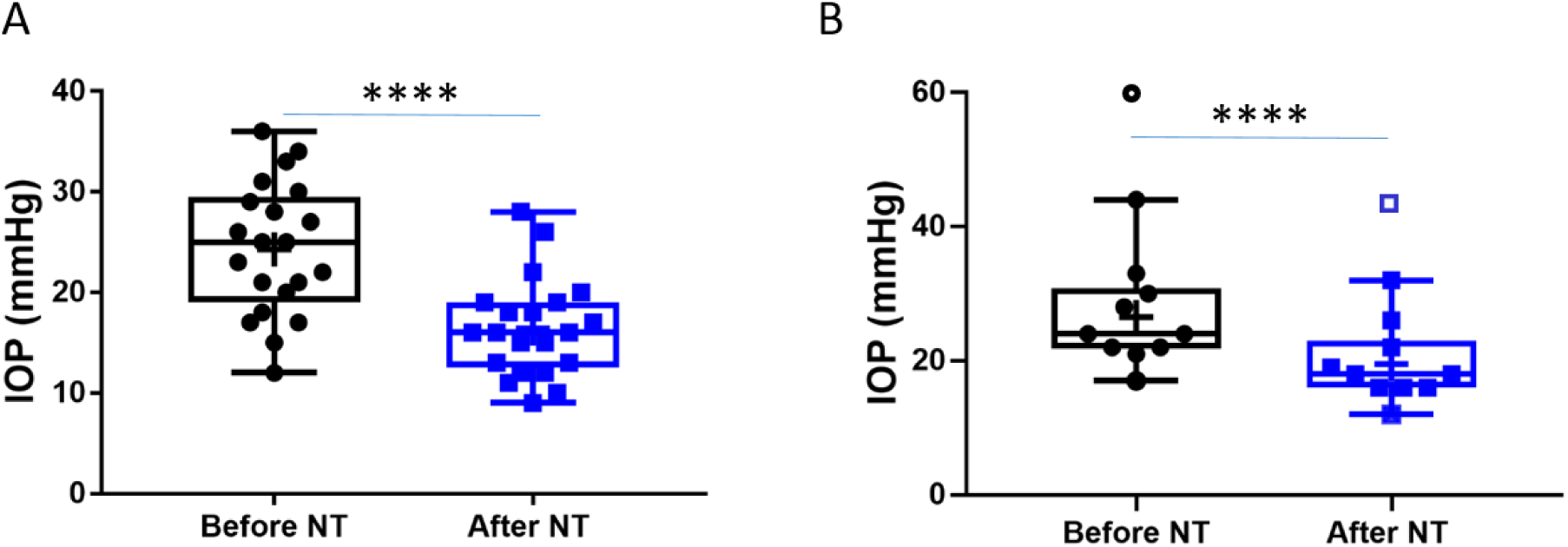
Netarsudil (NT) efficaciously lowered intraocular pressure (IOP) in patients with steroid-induced elevated IOP poorly controlled with standard glaucoma medications. IOPs were measured by Goldmann applanation tonometry in patients who demonstrated OHT after steroid treatment for a variety of ocular conditions (Table 1). These individuals were initially treated with aqueous humor suppressants and/or prostaglandin analogues but showed persistent OHT, and were thus treated with NT. Panel A shows IOPs of patient cohort 1 (n=21 eyes of 19 patients) and Panel B shows IOPs of patient cohort 2 (n=11 eyes of 8 patients). “Before NT” indicates IOPs measured before initiating NT in these patients, and “After NT” shows IOP after daily treatment with NT for one month or less. The central line in box and whisker represents the median, the top and bottom edges are 25th and 75th percentiles, the whiskers extend to the most extreme data points and “+” indicates the mean. Empty symbols were statistically determined to be outliers of data sets. ****p<0.0001.

To evaluate NT efficacy in a second cohort of patients, a retrospective chart review was conducted on all patients seen by an experienced glaucoma specialist (JRS) over a one-month period at two geographic locations. This cohort included patients with a single diagnosis of steroid-induced glaucoma that was uncontrolled on standard glaucoma medications, and who subsequently received NT. We identified 11 eyes from 8 patients, with an average age of 58.4 years (Table 1, Cohort 2). The IOP in these eyes prior to NT treatment was similar to the first cohort at 26.5 ± 7.7 mmHg (mean ± SD). As well, NT significantly lowered IOP in this second cohort by an average of 6.0 mmHg (p=0.0003, Figure 1B), leading to an IOP of 19.5 ± 5.8 mmHg (mean ± SD). We conclude that NT significantly lowers IOP in steroid glaucoma patients refractory to conventional anti-ocular hypertensive medications.

### Increase in outflow facility by NT effectively prevents and reverses steroid-induced OHT in a mouse model of human disease

To better understand the mechanism of NT’s IOP-lowering effect in patients, we carried out studies utilizing our established steroid-induced OHT mouse model (Li, Lee et al. 2019). It is known that daily treatment with NT significantly decreases IOP in naïve mouse eyes by improving conventional outflow function (Li, Mukherjee et al. 2016). Thus, we studied the efficacy of NT treatment in mice, focusing specifically on NT’s effect on conventional aqueous outflow dynamics and TM structure and function. Our experimental studies were designed to address two clinically important questions in this mouse model: *(1) Can NT prevent steroid-induced OHT?* and *(2) Can NT reverse steroid-induced OHT?*

To test prevention, mice were treated unilaterally with either NT or placebo (PL) starting one day *prior* to bilateral delivery of dexamethasone (DEX)-loaded nanoparticles (NPs), with NT or PL treatment continuing for the 4-week duration of DEX-NP exposure. Baseline IOPs in both NT and PL groups were similar (19.4 ± 0.4 and 19.6 ± 1.1 mmHg, respectively, p = 0.60). One day after NT or PL treatment, but before DEX treatment, IOP was 20.1 ± 0.9 mmHg in PL-treated eyes and significantly lower (16.1 ± 1.4 mmHg) in NT-treated eyes (p = 0.0009, Figure 2A). IOPs were followed for 4 weeks, and average IOP elevation in PL-treated eyes was 5.94 ± 0.57 mmHg, while IOP in NT-treated eyes was significantly lower than IOP in PL eyes (p < 0.0001; Figure 2B), returning essentially to baseline levels (average IOP elevation compared to baseline of 0.23 ± 0.45 mmHg, p = 0.71). Outflow facility, the primary determinant of IOP, was 84% greater in NT eyes compared to PL (4.87 ± 1.09 vs. 2.64 ± 0.44 nl/min/mmHg, p = 0.08, Figure 2C).

**Figure 2.**
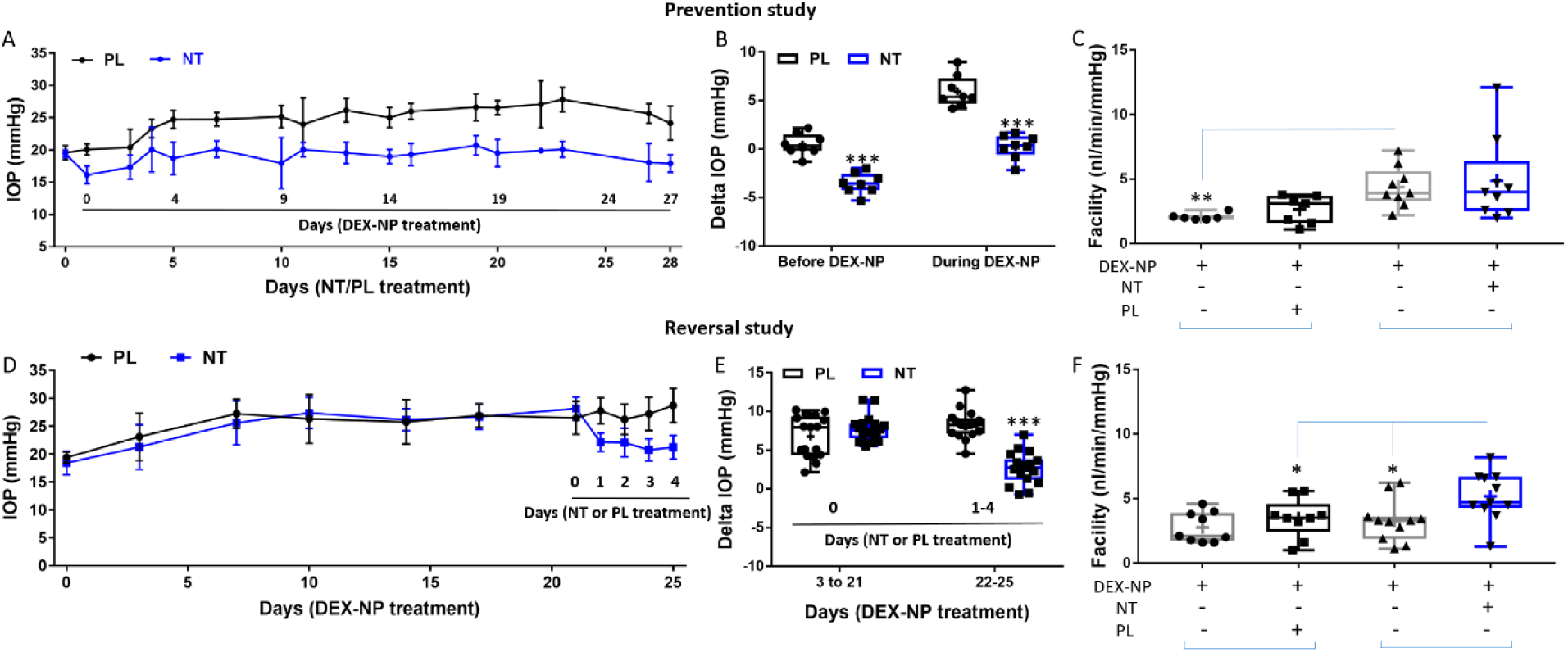
Netarsudil (NT) prevented and reversed steroid-induced ocular hypertension by improving outflow function. **(A)** In the prevention study, IOP was measured in two groups of age- and gender-matched wild-type C57BL/6 mice receiving NT or placebo (PL) unilaterally. The day after NT/PL application was started, dexamethasone-loaded nanoparticles (DEX-NPs) were delivered bilaterally into the periocular space to release DEX and create steroid ocular hypertension. We continued to apply NT or PL 2-3 times/week to the same eye and DEX-NPs 1-2 times/week bilaterally for 4 weeks The data are displayed as mean ± SD (n = 8 for each group). **(B)** We summarize the data from panel A by showing the average IOP elevation over baseline (“Delta IOP”) following NT or PL treatment in the presence of continuous DEX-NP delivery. “Before DEX-NP” refers to IOP elevations at 1 day post-NT or -PL treatment but before delivery of DEX-NPs, and “During DEX_NP” refers to IOP elevations averaged over 4-28 days of NT or PL treatment, i.e. over 3-27 days of DEX-NP delivery. ***, p < 0.001 comparing NT vs. PL groups. **(C)** After 4 weeks of PL or NT treatment, outflow facility was measured in freshly enucleated eyes (p = 0.08 for comparison of DEX-NP+PL vs. DEX-NP+NT, n = 7-9). Brackets indicate paired eyes. **(D)** In the second (reversal) study, we found that NT rapidly reversed three weeks of DEX-NP-induced ocular hypertension. IOP was measured in two groups of age- and gender-matched mice receiving DEX-NPs bilaterally 1-2 times/week for 4 weeks. During the last week of DEX-NP delivery, NT or PL was administered unilaterally once/day for 4 consecutive days. The data show mean ± SD (n = 17 for PL group and n = 19 for NT group). **(E)** We summarize the data from panel C by showing average IOP elevations above baseline (“Delta IOP”) for DEX-NP-treated eyes in both groups prior to NT or PL treatment (left side) or averaged over 1-4 days of NT or PL treatment (right side). ***, p < 0.001 comparing NT vs. PL groups. (**F**) On day 5, both eyes were enucleated and outflow facility was measured (* p = 0.038, n = 9 for DEX-DP+PL and n = 11 for DEX-NP+NT group). Brackets indicate paired eyes. See Fig 1 for interpretation of box and whisker plots.

Motivated by these findings, we next tested NT’s ability to reverse steroid-induced OHT. We delivered DEX-NPs bilaterally for 3 weeks to mice *before* initiating unilateral NT or PL therapy (once/day for 4 consecutive days). The baseline IOPs before DEX treatment in both NT and PL groups were similar (18.5 ± 1.76 vs. 19.2 ± 1.28 mmHg, respectively; p = 0.21). After 3 days of DEX-NP administration, IOP was significantly elevated in both groups (Figure 2D). After three weeks of DEX-NP treatment, average IOP elevation (days 3-21) compared to baseline in PL and NT cohorts was similar (6.77 ± 0.67 mmHg vs. 7.74 ± 0.384 mmHg, respectively, p = 0.45, Figure 2E). Commencement of NT treatment resulted in rapid IOP lowering (within 1 day) followed by a continued decrease. After 4 days of treatment, the change in IOP from baseline in PL-treated eyes was 8.19 ± 0.46 versus 2.69 ± 0.47 mmHg for NT-treated eyes (p < 10^−4^). In fact, NT reversed steroid-induced OHT, returning IOP to near baseline levels (18.5 ± 1.76 vs. 21.4 ± 1.30 mmHg). NT increased outflow facility by 33% compared to PL (5.18 ± 0.57 vs. 3.48 ± 0.51 nl/min/mmHg, p = 0.038, Figure 2F) and increased outflow by 37% compared to contralateral eyes (3.27 ± 0.49 nl/min/mmHg, p = 0.025). Similar to the prevention study, these measured facility differences can mathematically account for ∼100% of the observed IOP differences between NT- and PL-treated eyes. We conclude that NT’s IOP lowering effect in this steroid glaucoma model can be largely explained by increased outflow facility.

### Reversal of steroid-induced trabecular meshwork stiffening by NT

Mechanical stiffness of the TM, a key tissue in the conventional outflow pathway, was shown to be increased by GC treatment, negatively correlating with outflow facility (Wang, Li et al. 2018). We also previously showed that GC treatment decreased the tendency of the SC lumen to collapse at elevated IOPs (Li, Lee et al. 2019), an effect that appeared to be mediated by changes in TM stiffness. Since rho kinase inhibitors such as NT decrease cellular contractility and act as anti-fibrotic agents (Lin, Sherman et al. 2018), we hypothesized that NT would reverse steroid-induced conventional outflow tissue stiffening and lead to more SC collapse at elevated IOPs. To test this hypothesis, we used our reversal protocol as above. On day 5 after the last NT/PL treatment, mice were anesthetized and secured on a custom imaging platform (Li, Farsiu et al. 2014, Li, Farsiu et al. 2014, Boussommier-Calleja, Li et al. 2015, Li, Mukherjee et al. 2016, Li, Lee et al. 2019). The anterior chamber of the NT- or PL-treated eye was cannulated with a single needle connected to a fluid reservoir, allowing IOP to be controlled. The conventional outflow tissues were imaged using OCT as IOP was clamped at 5 different levels. With increasing IOP, the SC lumen became smaller in both NT- and PL-treated eyes, but to different extents. In PL-treated eyes, the SC lumen was still patent at an IOP of 20 mmHg, while the SC lumen in NT-treated eyes was almost completely collapsed. In fact, the SC luminal cross-sectional areas differed significantly between NT- and PL-treated groups at each pressure level (p = 0.0024, Figure 3 and Figure S1).

**Figure 3.**
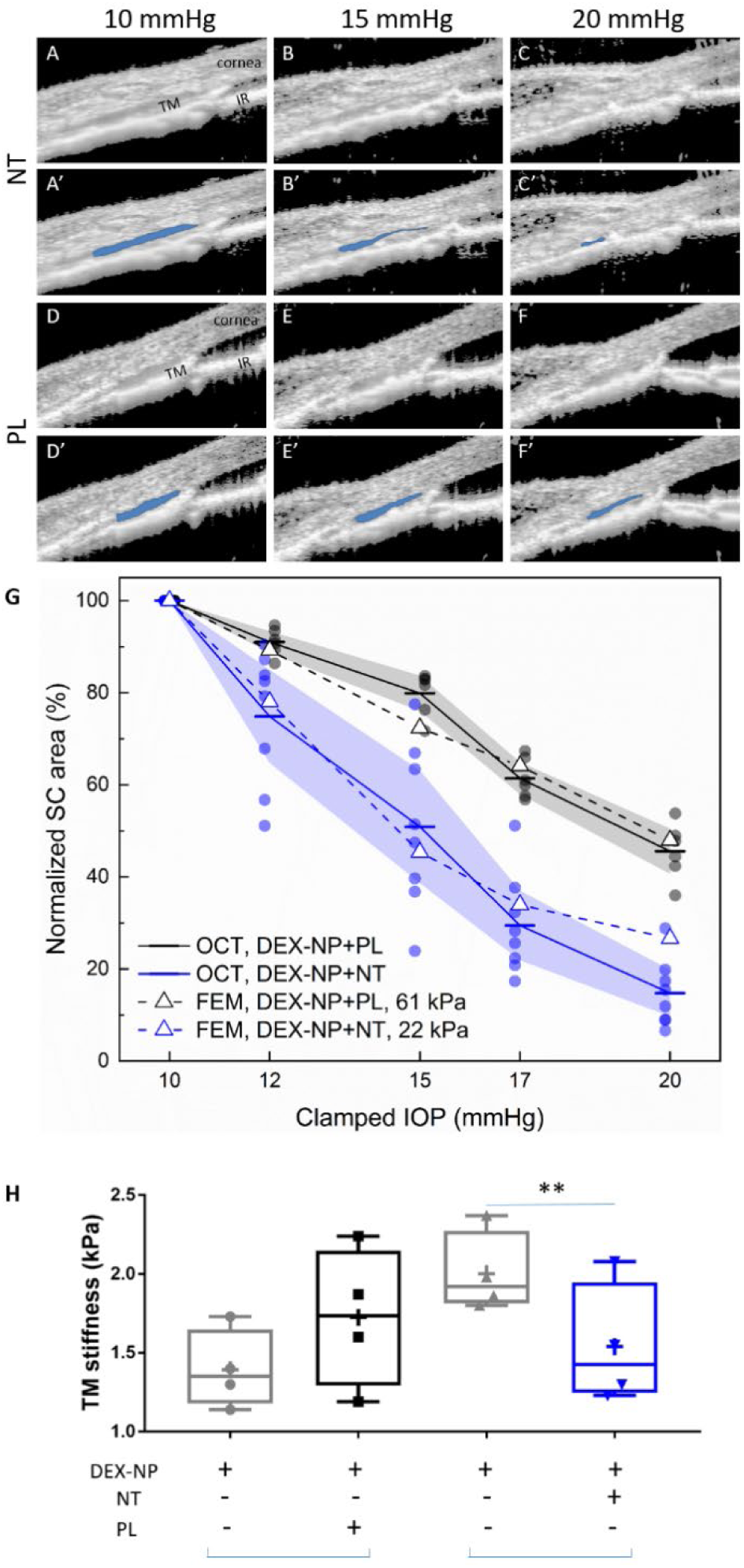
Netarsudil (NT) reduced steroid-induced TM stiffening visualized in living mice by SD-OCT and estimated by inverse finite element modeling and measured directly by atomic force microscopy. Wild-type C57BL/6 mice received dexamethasone-loaded nanoparticles (DEX-NPs) bilaterally for 4 weeks. During the last week, mice received either NT or placebo (PL) unilaterally for 4 consecutive days. **(A-F)** On day 5, living mouse eyes were cannulated to control IOP and were subjected to sequentially increasing pressure steps (10-20 mmHg) while imaging conventional outflow tissues using OCT. Images were analyzed and SC lumen semi-automatically segmented (in blue) using custom SchlemmSeg software **(A’-F’)**.The increased tendency towards SC collapse in NT-treated vs. PL-treated eyes is evident. **(G)** Quantitative comparison of SC lumen areas (blue regions in panels A’-F’) in both NT and PL treatment group at 5 clamped IOPs (10, 12, 15, 17 and 20 mmHg). The plotted quantity is relative SC area (normalized to value at 10 mmHg), and the data show an increased tendency toward SC collapse in NT-treated eyes compared to PL-treated eyes. The dots indicate individual eyes, and bars represent mean values for each IOP. Shaded regions indicate 95% confidence intervals. N = 6 for PL and n = 8 for NT treatment groups. We conducted inverse finite element modeling (iFEM) to structurally analyze the response of anterior segment tissues to varying IOP levels, mimicking the experimental measurements. Dashed lines show results of the iFEM analysis for least-squares best fit TM stiffness values, yielding estimated stiffnesses of 61 kPa for PL-treated eyes and 22 kPa for NT-treated eyes. Abbreviations: IR = iris. **(H)** At day 5, eyes were collected and processed for atomic force measurements of TM stiffness (Young’s modulus). We observed that netarsudil softened the TM vs. control eyes. Brackets indicate paired eyes. We point out that the magnitudes of the AFM and OCT/iFEM measurements of TM stiffness differ due to the well-known dependence of soft tissue stiffness on loading mode (compression by AFM vs. tension by SC lumen pressurization) (Ethier and Simmons 2007), as discussed in more detail in (Wang, Johnstone et al. 2017). An average of 135 measurements were made from 3 quadrants on 4 biological samples for both cohorts, p =0.007. Blue brackets indicate paired eyes. See Figure 2 for interpretation of box and whisker plots.

We next quantified NT effects on TM stiffness using inverse finite element modeling (iFEM) (Li, Lee et al. 2019) based on the OCT images acquired in vivo. This iFEM procedure, allowing us to deduce TM stiffness based on structural analysis of SC collapse and associated TM deformation, was performed on OCT images from both NT- and PL-treated groups. We deduced TM tissue stiffness values of 22 kPa in the NT-treated group vs. 61 kPa in the PL-treated group (Figure 3G). The latter value is consistent with the TM stiffness we previously measured in DEX-NP treated eyes of 69 kPa (Li, Lee et al. 2019). Remarkably, the TM stiffness we estimated in NT-treated eyes was close to the value of 29 kPa that we previously determined in naïve eyes (Li, Lee et al. 2019) (Figure S3). Thus, it appears that NT returns TM stiffness to control levels after only 4 days of administration in eyes made hypertensive by long-term GC administration.

It was desirable to obtain an independent and direct measurement of TM stiffness in a cohort of NT-treated mouse eyes. For this purpose, we used atomic force microscopy (Wang, Read et al. 2017, Wang, Li et al. 2018, Li, Lee et al. 2019). Using the reversal treatment protocol, we observed that NT significantly reduced TM tissue stiffness by 23% compared to contralateral eyes (1.54 ± 0.19 vs. 2.00 ± 0.13 kPa, p = 0.007, Figure 3H). In contrast, PL did not affect TM tissue stiffness compared to contralateral eyes (1.39 ± 0.12 vs. 1.73 ± 0.20 kPa, p = 0.39). When stiffness measurements of PL and NT groups were normalized to untreated contralateral eyes, a trend towards NT-induced softening was observed but was not significant due to small sample size (p=0.15).

### Reversal of GC-induced fibrotic alterations in conventional outflow tissues by NT

We next investigated NT effects on fibrotic and morphological changes in TM tissues. At the light microscopic level, we observed no gross morphological changes in conventional outflow tissues in NT-versus PL-treated steroid-induced OHT eyes (Figures 4A and 4B). In contrast, we observed that NT treatment significantly reduced the expression of alpha smooth muscle actin (αSMA; Figures 4C and 4D) and fibronectin (FN; Figures 4E and 4F), two fibrotic indicators known to be elevated in conventional outflow tissues after GC treatment. In fact, NT-mediated down-regulation led to αSMA and fibronectin levels similar to those observed previously in eyes treated with PBS-loaded (sham) nanoparticles (Li, Lee et al. 2019). Moreover, when examined at the electron microscopic level, we observed three major effects of NT treatment on steroid-induced OHT eyes. The first was a significant reduction in the amount and density of basal lamina materials underlying the inner wall of SC (n=7, p=0.02, Figure 5). These deposits were scored on a semi-quantitative scale; confirming qualitative impressions of NT treatment (Figure 5D). The second was an apparent increase in the number of “open spaces” in the TM of NT-treated eyes, including spaces between lamellar beams and in the JCT region. Lastly, we observed a change in the morphology of inner wall cells, with increased “ruffling” in the presence of NT.

**Figure 4.**
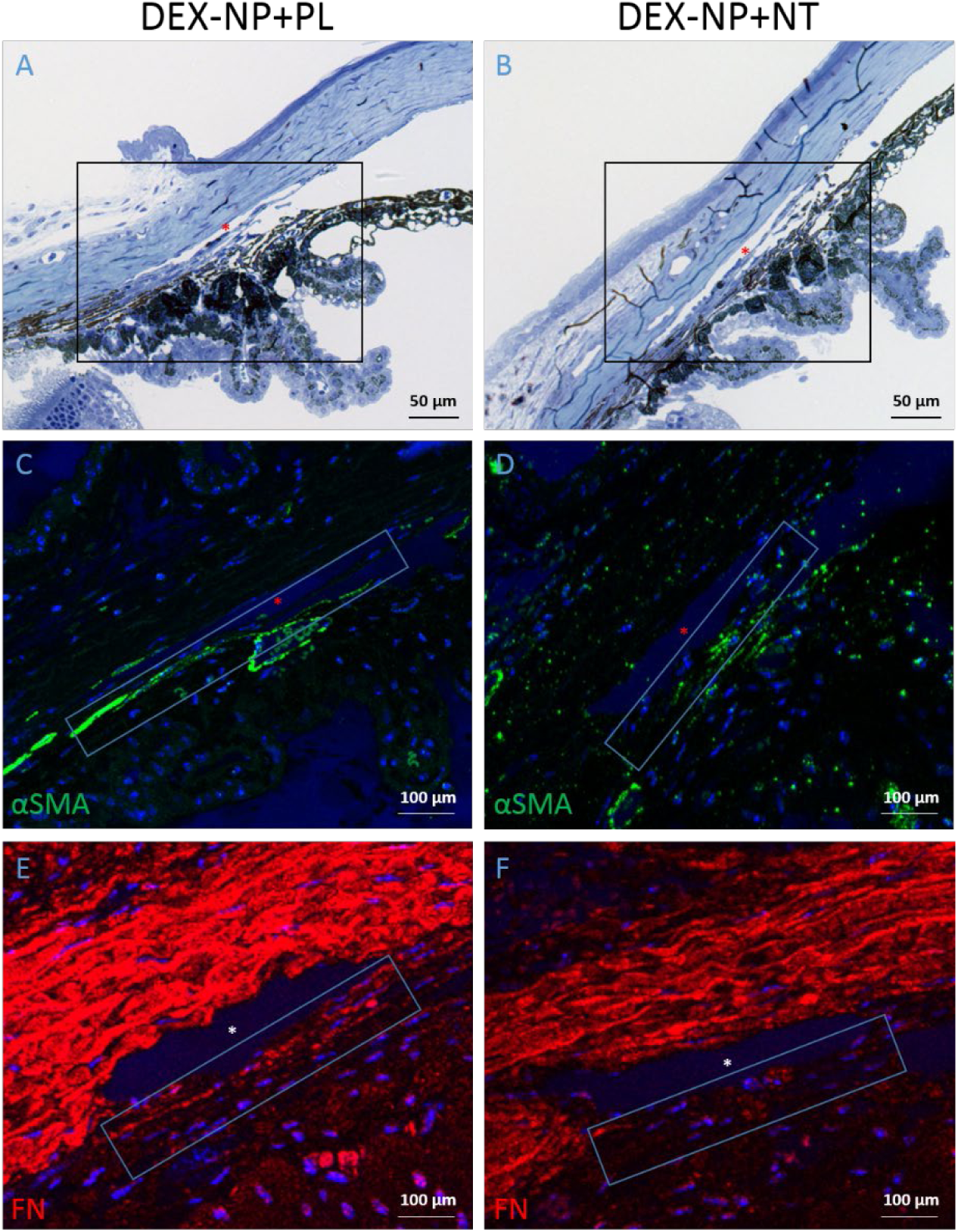
Netarsudil (NT) decreased fibrotic markers in ocular hypertensive mouse eyes. Two groups of age- and gender-matched wild-type C57BL/6 mice had dexamethasone-loaded nanoparticles (DEX-NPs) delivered bilaterally 1-2 times/week for 4 weeks to create ocular hypertension. During the last week of DEX-NP exposure, mice were split into two subgroups and treated unilaterally with either NT or placebo (PL) once/day for 4 consecutive days. At day 5, eyes were collected and processed for histological analysis. **(A and B)** Representative sagittal sections of iridocorneal tissues visualized by light microscopy after methylene blue staining, showing normal gross morphology in NT-treated eyes. Boxes in A and B indicate areas of interest displayed in C,E and D,F, respectively. **(C-F)** PL- or NT-treated ocular hypertensive eyes were sectioned and iridocorneal tissues were probed with antibodies recognizing α-smooth muscle actin (αSMA; C-D) or fibronectin (FN; E-F). Identical confocal settings were used for both NT and PL treatment groups. Data shown are representative of samples from 5 images from 4 mice treated with NT and 7 images of 5 mice treated with PL. Nuclei were counterstained with DAPI in (C-F). Asterisks indicate SC lumen. The images are representative of at least 4 animals per group.

**Figure 5.**
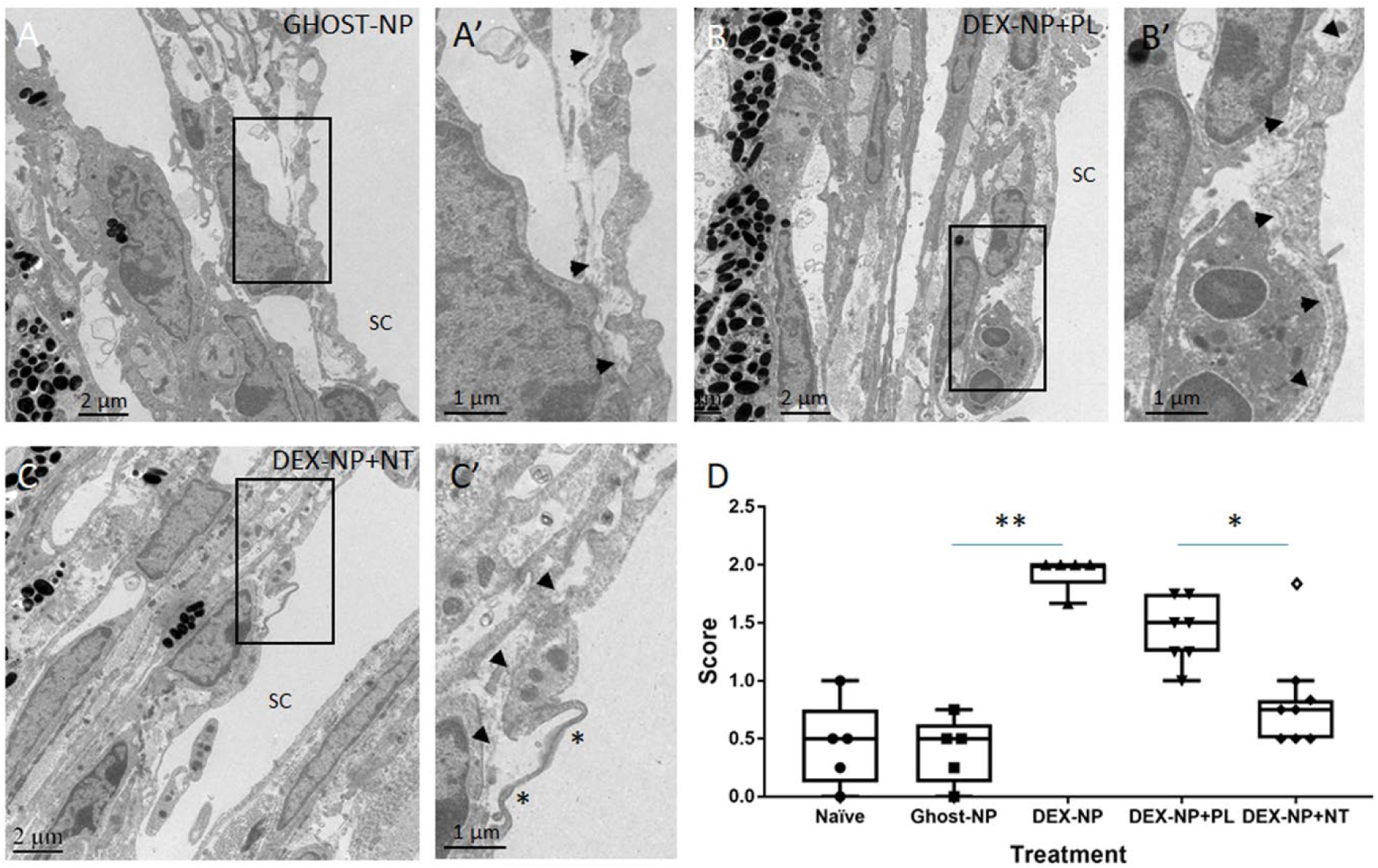
Netarsudil (NT) partially normalized ultrastructure of conventional outflow tissues in ocular hypertensive eyes. Two groups of age- and gender-matched wild-type C57BL/6 mice were bilaterally treated with dexamethasone-loaded nanoparticles (DEX-NPs) 1-2 times/week for 4 weeks. During the last week of DEX-NPs exposure, one group of mice was unilaterally treated with NT and another group with placebo (PL) once/day for 4 consecutive days. At day 5, eyes were collected and fixed with 4% PFA plus 1% glutaraldehyde in PBS for 1-3 days at 4°C. The anterior segments were embedded in Epon, sectioned, stained with uranyl acetate/lead citrate, and examined with a JEM-1400 electron microscope. We show representative images from **(A)** 5 control eyes (exposed to nanoparticles only without DEX [Ghost-NP]), 5 DEX-NP treated eyes, including **(B)** 7 PL-treated eyes and **(C)** 8 NT-treated eyes. These images correspond to 10, 13, 18, 12 and 8 images in each group that were graded. (A’-C’) show enlarged areas indicated by boxes in (A-C). Arrowheads point to basement membrane materials underlying SC inner wall endothelial cells. Asterisk indicates a SC membrane ruffle. **(D)** Results from scoring basal lamina appearance underlying SC endothelial cells. Images in panel A and C were scored as “0” (normal appearance), while images in panel B were scored as “2” (continuous and extensive basal lamina material). **, p <0.01 comparing Ghost-NP vs. DEX-NP; *, p < 0.05 comparing DEX-NP+PL vs. DEX-NP+NT. Empty symbol was statistically determined to be outlier of DEX-NP+NT data set.

## Discussion

The major finding of the current study was that NT effectively reversed steroid-induced OHT in two different cohorts of patients who were refractory to standard glaucoma medications. This clinical observation was mechanistically studied in a reliable, established mouse model of steroid-induced glaucoma (Li, Lee et al. 2019). As in patients, NT rapidly rescued long-term steroid-induced OHT in mice, and also effectively prevented the onset of steroid-induced OHT. NT-mediated rescue of OHT in mice was accompanied by restoration of normal outflow facility and conventional outflow tissue stiffness, as well as significant morphological alterations in the TM. Calculations suggest that the IOP effect seen in NT-treated mice was largely or entirely explainable by changes in outflow facility, which together with our direct and indirect measurements of TM stiffness, strongly implicate the TM as the tissue mediating the effects of NT on IOP. Significantly, this is the first demonstration of efficacious prevention and rescue from steroid-induced OHT by a “TM-active” FDA-approved glaucoma medication and strongly suggests that ROCKi compounds have effective anti-fibrotic activity in the trabecular meshwork.

Our results extend findings from an earlier report showing that a ROCKi lowered IOP in a transgenic mouse locally overexpressing CTGF (Junglas, Kuespert et al. 2012), to a clinically-relevant disease context by demonstrating NT activity in the conventional outflow pathway. Importantly, two significant phenotypic changes were observed in NT-treated eyes with OHT. The first was a rapid reversal of IOP elevation. The second was a partial clearance of accumulated ECM materials, namely fibronectin and basal lamina material underlying the inner wall of SC. Both cellular contractility changes and ECM changes likely contribute to the observed decrease of TM tissue stiffness by NT. In addition to NT-mediated changes in ECM turnover, the observed changes in ECM composition and amount may also be due to NT-mediated opening of flow pathways and consequential removal of ECM. In any case, having a therapy for steroid glaucoma that reverses the vicious cycle of fibrosis by restoring function to a diseased tissue offers a previously unavailable potential benefit for patients (figure 6).

**Figure 6:**
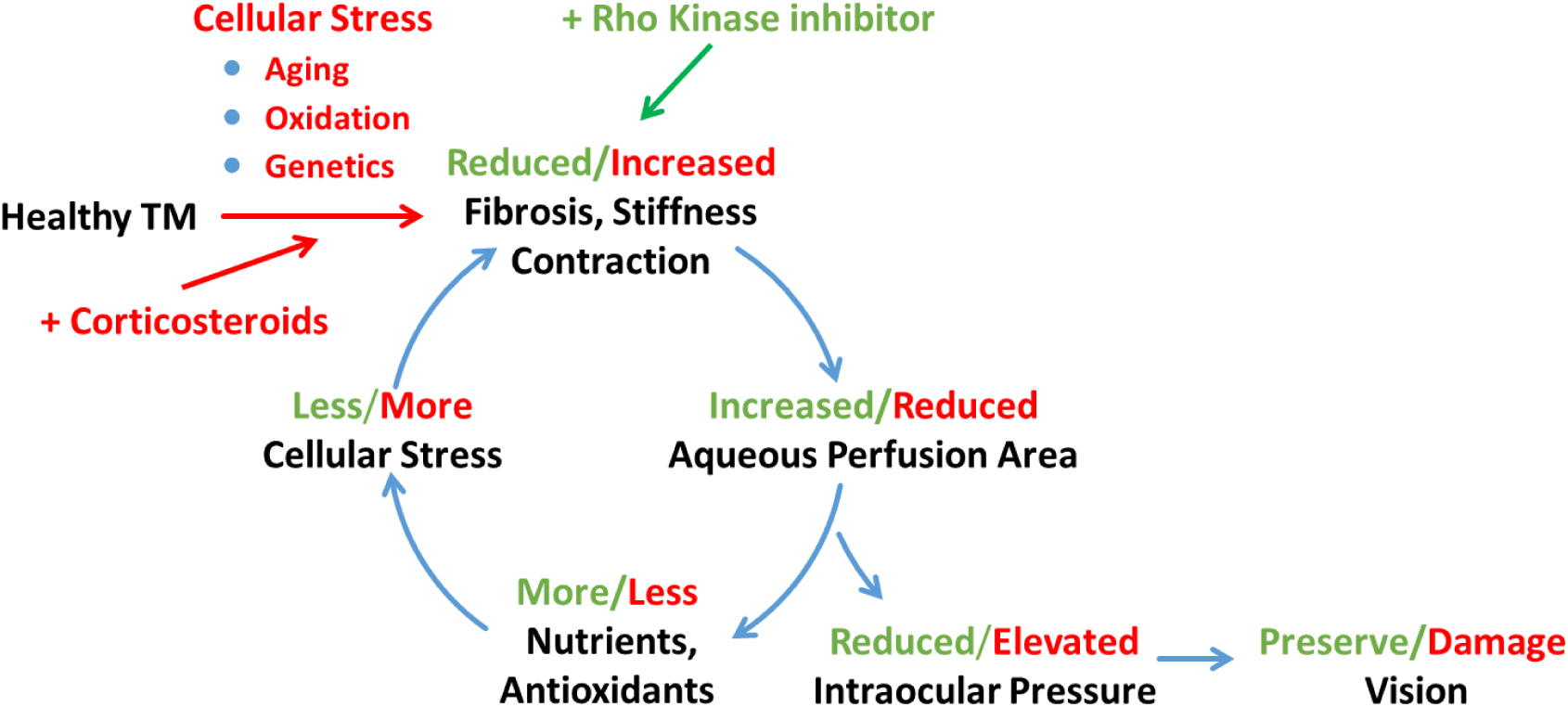
Schematic representation of a feed-forward model of fibrotic disease in the conventional outflow pathway responsible for ocular hypertension, incorporating a number of pathophysiological aspects of ocular hypertension (Stamer and Acott 2012, Stamer, Braakman et al. 2015). The broader literature suggests that this feed-forward loop can be triggered by multiple factors, including aging, oxidative stress or genetic predisposition. In this work we triggered this loop by corticosteroid administration and restored tissue function through rho kinase inhibition. TM: trabecular meshwork.

Evidence shows that the biomechanical properties of TM tissue play an important role in the regulation of outflow function, and thus IOP (Wang, Johnstone et al. 2017, Wang, Li et al. 2018). For example, alterations in the biomechanical properties (i.e., stiffening) of the TM are observed in glaucomatous human donor eyes, which were hypothesized to be associated with dysregulation of the ECM (Lutjen-Drecoll, Shimizu et al. 1986, Yue 1996, Tamm and Fuchshofer 2007, Keller, Aga et al. 2009, Tektas and Lutjen-Drecoll 2009, Last, Pan et al. 2011). TM stiffening and elevated IOP was also observed in animal models of steroid-induced OHT (Raghunathan, Morgan et al. 2015, Li, Lee et al. 2019). In a recent study, we showed that TM tissue stiffness is associated with increased outflow resistance in steroid-induced OHT mouse eyes (Wang, Li et al. 2018). We consider it remarkable that NT appears to have restored normal TM biomechanical properties in our steroid-induced glaucoma model after only 4 days of treatment, and that this restoration was accompanied by significant morphological changes in the extracellular matrix (ECM) of the TM. These in vivo observations in a relevant disease model powerfully confirm the importance of the rho-kinase pathway, and more generally the importance of cellular biomechanical tension as mediated through actomyosin cytoskeletal tension, on ECM synthesis, assembly and degradation (Rao, Deng et al. 2001, Nakajima, Nakajima et al. 2005, Pattabiraman and Rao 2010). The rapid time scale of the observed ECM changes is consistent with observations from post mortem human eye preparations (Keller, Aga et al. 2009). The time course of structural changes in the TM was also consistent with the observed rapid (within 24 hours) pharmacodynamics of NT on IOP, completely reversing 1 and 3 months of OHT due to DEX treatment (Figure 2 and Figure S4, respectively). This time course is consistent with known effects of rho kinase inhibitors (McMurtry, Abe et al. 2010, Lin, Sherman et al. 2018).

More generally, these data emphasize the potent, anti-fibrotic properties of ROCKi in the conventional outflow pathway, consistent with a large body of literature in other tissues. Many mechanisms have been proposed for the systemically-delivered anti-fibrotic effects of ROCKi, including inhibition of monocyte differentiation in a murine model of ischaemic/reperfusion cardiomyopathy (Haudek, Gupta et al. 2009), and macrophage infiltration of injured tissues in mouse models of multiple fibrotic diseases, including renal tubulointerstitial fibrosis, diabetic nephropathy, renal allograft rejection, peritoneal fibrosis, and atherosclerosis (Wu, Xu et al. 2009, Knipe, Tager et al. 2015). In view of the eye’s immune-privileged status and local delivery of NT in the present study, it seems likely that resident cells of the conventional outflow pathway primarily mediate NT effects. In other tissues, RKIs inhibit TGF-β1-induced activation of p38 MAPK (Holvoet, Devriese et al. 2017), activation of myocardin-related transcription factors (MRTFs), known to be master regulators of epithelial-mesenchymal transition (Gasparics and Sebe 2018), and reduced activation of NF-κB (Segain, Raingeard de la Bletiere et al. 2003). Importantly, downstream targets of MRTFs include CTGF and YAP/TAZ, and TGFβ-mediated signaling through p38 and NF-κB (Raghunathan, Morgan et al. 2013, Braunger, Fuchshofer et al. 2015, Inoue-Mochita, Inoue et al. 2015, Tamm, Braunger et al. 2015, Montecchi-Palmer, Bermudez et al. 2017, Hernandez, Roberts et al. 2020); all of which participate in the regulation of trabecular meshwork contractility, mechanotransduction and intraocular pressure homeostasis. Thus, our DEX-induced ocular hypertension model represents a powerful tool for further interrogation of these pathways in the context of an immune-privileged environment (Figure 6).

We used two complementary methods to estimate conventional outflow tissue stiffness, and each has advantages and disadvantages. AFM was used to directly measure the compressive mechanical properties of the conventional outflow tissues in mouse eyes at different locations of the TM and around the eye with high resolution (Tao, Lindsay et al. 1992, E, Heinz et al. 1998, Matzke, Jacobson et al. 2001, Wang, Read et al. 2017). However, AFM measurements of stiffness could only be conducted ex vivo on “dead” tissues using a cryosectioning technique coupled with AFM. Thus, these measurements likely reflect the stiffness of the ECM, not cells (Wang, Read et al. 2017). In contrast, tensile stiffness estimates of conventional outflow tissues were derived from changes in area of SC lumen in living mouse eyes exposed to sequential IOP challenges. TM behavior was captured using SD-OCT, and stiffness was then quantified by inverse FEM as described previously (Li, Lee et al. 2019). The disadvantage with OCT is that eyes are imaged at only one location. Regardless, both methods showed consistent results, i.e. that NT significantly reduced steroid-induced conventional outflow tissue stiffness. While desirable, technical concerns prevented OCT, AFM and outflow facility measurements in the same eye.

Before the approval of NT in 2017, four classes of glaucoma drugs (beta-adrenergic receptor antagonists, alpha-adrenergic agonists, carbonic anhydrase inhibitors and/or prostaglandin analogues) were used to treat steroid-induced OHT. Unfortunately, none of these drugs directly target tissues responsible for homeostatic regulation of IOP in healthy eyes, nor for the dysregulation of IOP in steroid-induced glaucoma. Instead, these drugs either decrease aqueous humor formation or divert aqueous humor from the conventional outflow pathway by increasing unconventional outflow drainage (Bucolo, Platania et al. 2015, Schmidl, Schmetterer et al. 2015). Despite modest effects on conventional outflow (Brubaker, Schoff et al. 2001, Wan, Woodward et al. 2007, Bahler, Howell et al. 2008), prostaglandins are not often used to treat steroid-induced OHT due to concern about their effects on the blood-retinal barrier in vulnerable retinas (Table 1, (Moroi, Gottfredsdottir et al. 1999)). The recent approval of NT changed the paradigm of targeting glaucomatous disease, being the first agent to selectively target and modify conventional outflow morphology and function (Li, Mukherjee et al. 2016, Ren, Li et al. 2016). However, the current standard of care is for patients to be first treated with glaucoma medications that do not target the conventional outflow pathway, with NT being used only when these drugs are ineffective at IOP lowering. Thus, in two independent patient populations we found 27 patients that had steroid-induced OHT and were treated with NT. Despite being first treated with multiple first- and second-line glaucoma medications (mean of 2.7 and 2.4), NT lowered IOP by an average of 7.9 mmHg in the first cohort and 6.0 mmHg in the second. These observations are consistent with mechanistic data from our mouse model, supporting the concept that the TM is the location of pathology in steroid-induced OHT, and emphasizing the importance of targeting the conventional outflow pathway in this well-recognized condition.

Steroid-induced OHT in mice was generated using nanotechnology (Agrahari, Li et al. 2017), resulting in a model that closely matched steroid-induced observations in the human condition (decreased outflow facility, increased accumulation of ECM in the TM, and elevated IOP) (Johnson, Bradley et al. 1990, Johnson and Knepper 1994, Johnson, Gottanka et al. 1997, Clark, Steely et al. 2001). The present study further highlights the utility of this mouse model of steroid-induced glaucoma and its translational relevance to the human clinical condition. The first, and most direct, outcome of this mouse model in the present work is motivation for considering the use of NT to prevent or treat OHT caused by GCs, their most challenging adverse event. This is particularly pertinent in view of the high incidence of OHT in patients receiving intravitreal GC-delivery implants to treat uveitis or macular edema. This motivation is reinforced by our data showing that human steroid-induced OHT patients who are refractory to standard glaucoma medications respond very well to NT. This retrospective study encourages a prospective study testing NT as a first-line drug for steroid-induced ocular hypertension. The second outcome of our study is to advance and validate methods for non-contact, non-invasive estimation of conventional outflow function/health using OCT and iFEM. Our results confirm that this technology has the resolution to detect changes in conventional outflow dysfunction due to GC treatment and restoration of function after drug treatment. In clinical practice, we suggest that this technology may have potential utility in staging glaucoma status and in monitoring treatment.

## Materials and Methods

### Study Design

Experiments were designed to test the hypothesis that dysregulation of conventional outflow function caused by glucocorticoids can be mitigated by NT treatment. The hypothesis was tested using a retrospective review of patients who were refractory to standard glaucoma treatments, and in a validated mouse model of steroid-induced ocular hypertension that we developed (Li, Lee et al. 2019).

Data from humans were obtained using an unbiased retrospective analysis of electronic medical records of all patients seen by ophthalmologists at Duke Eye Center and from a retrospective chart review of two private clinical practices. All steroid-induced glaucoma patients treated with NT were included if not complicated with secondary diagnosis as detailed below. IOPs from one patient were excluded from the data set because they were greater than 1.5 times a quartile different from 75^th^ percentile.

Mouse studies were based on sustained peri-ocular delivery of steroid to induce ocular hypertension (Li, Lee et al. 2019). Cohort sizes were determined using power analysis of data generated previously with this model (Li, Lee et al. 2019) and treatment effects with NT (Li, Mukherjee et al. 2016). Eyes of mice were randomized to receive either topical NT or PL, and experimentalists were masked to treatment type. Primary endpoints (IOP, outflow facility, Young’s modulus of TM, outflow tissue behavior visualized by OCT following IOP challenges, outflow tissue morphology by TEM and fibrotic marker expression by IHC) were established prior to start of experiments and all data were included in the analyses.

### Patient Ascertainment, IOP Measurement and Treatment

We identified two cohorts of patients with steroid-induced ocular hypertension who had been treated with NT. The first cohort was drawn from patients seen at the Duke Eye Center, using IRB-approved access to patient data via the EPIC electronic medical record system using SlicerDicer software. Search criteria included “netarsudil” and associated ICD-10 codes for “steroid responder” and “steroid glaucoma” (H40.041, H40.042, H40.043, T38.0×5A, H40.60×0, H40.61×0, H40.62×0, and H40.63×0) between 1/1/2018 and 3/1/2020. Twenty-one eyes of nineteen patients were identified as having started NT due to steroid-associated intraocular pressure elevation. None of the patients were “tapered” from their steroid during the first month of treatment.

The second cohort was based on a comprehensive retrospective chart review of patients seen by Dr. John Samples at two different clinic sites: The Eye Clinic in Portland, Oregon and Olympia Eye Clinic in Olympia, Washington. Charts of all patients seen by Dr. Samples between November 3 to December 1, 2019 were reviewed. Patients were included in the study that: (i) had a diagnosis of steroid-induced glaucoma, (ii) were uncontrolled on standard glaucoma medications, (iii) were treated with NT, and (iv) had no confounding diagnoses (exfoliation glaucoma, pigmentary glaucoma, active neovascularization, narrow/closed angles, previous glaucoma surgery).

IOP in both studies was measured by Goldmann applanation tonometry before starting NT and then within one month after QD NT treatment.

### Animals

C57BL/6 (C57) mice (both males and females, ages from 3-6 months) were used in the current study. The animals were handled in accordance with approved protocols (A020-16-02 and A001-19-01, Institutional Animal Care and Use Committee of Duke University) and in compliance with the Association for Research in Vision and Ophthalmology (ARVO) Statement for the Use of Animals in Ophthalmic and Vision Research. The mice were purchased from the Jackson Laboratory (Bar Harbor, Maine, USA), bred/housed in clear cages and kept in housing rooms at 21°C with a 12h:12h light:dark cycle.

### Ocular hypertension animal model

Ocular hypertension in mice was created by injection of nanoparticles entrapping dexamethasone (DEX-NPs) into either the subconjunctival or periocular spaces as described in previous publications (Wang, Li et al. 2018, Li, Lee et al. 2019). Briefly, DEX-NPs were diluted in PBS to a final NP concentration of 50 µg/µl, vortexed for 10 min and then sonicated for 10 min. Mice were anesthetized with 100mg/10mg/kg of ketamine/xylazine. 20 µl per eye of the DEX-NP suspension (containing 1mg of NPs with ∼23 µg of DEX) was slowly injected bilaterally into either the superior or inferior subconjunctival or periocular spaces using a 30-gauge needle with a Hamilton glass microsyringe (50 µl volume; Hamilton Company, Reno, NV). After withdrawing the needle, Neomycin plus Polymyxin B Sulfate antibiotic ointment was applied to the eye and mice recovered on a warm pad. The injection was conducted 1-2 times/week for 3-4 weeks. For DEX-NP control treatment, nanoparticles without DEX (Ghost-NP) were used to treat mice in the same way.

### Drug treatments

NT and placebo eye drops were provided in de-identified dropper bottles by Aerie Pharmaceuticals, Inc. All mice received bilateral treatments of DEX-NP, but were randomized as to whether they were given NT or PL. In the prevention study, mice were treated unilaterally with 0.04% NT or placebo (PL) by subconjunctival injection of 10 µl of NT or PL 1 day prior to bilateral DEX-NP treatment as described above, followed by NT or PL delivery 2-3 times per week for the entire 4 week duration of DEX-NP delivery. In the reversal study, eyes received DEX-NPs bilaterally 1-2 times per week for 4 weeks. During the last week of DEX-NP delivery, mice were treated unilaterally with NT or PL by subconjunctival injection of 10µl NT or PL once/day for four days.

### Intraocular pressure measurements

Mice were anesthetized with ketamine (60 mg/kg) and xylazine (6 mg/kg). IOP was measured immediately upon cessation of movement (i.e., in light sleep) using rebound tonometry (TonoLab, Icare, Raleigh, NC) between 10am and 1pm (Li, Farsiu et al. 2014, Li, Farsiu et al. 2014, Meng, Nikolic-Paterson et al. 2016, Li, Torrejon et al. 2018, Li, Lee et al. 2019). Each recorded IOP was the average of six measurements, giving a total of 36 rebounds from the same eye per recorded IOP value. IOP measurements were conducted twice per week.

### Outflow facility measurements

Outflow facility was measured by a technician (IN) who was masked as to the treatment group using the iPerfusion system as described in detail previously (Li, Mukherjee et al. 2016, Li, Lee et al. 2019). Briefly, at the end of the 4 day NT/PL treatment period in the reversal study, and after 4 weeks of NT/PL treatment in the prevention study, mice were euthanized using isoflurane, and eyes were carefully enucleated and rapidly mounted on a stabilization platform located in the center of a perfusion chamber using a small amount of cyanoacrylate glue (Loctite, Westlake Ohio, USA). The perfusion chamber was filled with pre-warmed Dulbecco’s phosphate-buffered saline with added 5.5 mM D-glucose (DBG), submerging the eyes and regulating temperature at 35°C. Two glass microneedles, back filled with filtered DBG, were connected to the system. Using micromanipulators, one microneedle was inserted into each anterior chamber of paired eyes without contacting the irises. Both eyes were perfused at 9 mmHg for 30 min to allow acclimatization and stabilization, followed by perfusion at 9 sequential pressure steps of 4.5, 6, 7.5, 9, 10.5, 12, 15, 18 and 21 mmHg. Poor quality steps and subsequent pressure steps were eliminated using established criteria (Sherwood, Reina-Torres et al. 2016). Stable flow rate (Q) and pressure (P) averaged over 4 minutes at each pressure step were used for data analysis to compute outflow facility (Meng, Nikolic-Paterson et al. 2016, Sherwood, Reina-Torres et al. 2016, Li, Torrejon et al. 2018).

### Calculations Relating Outflow Facility Changes and IOP Changes

We asked whether measured IOP differences between NT and PL cohorts were quantitatively consistent with measured differences in outflow facility between these cohorts. For this purpose, we assumed that unconventional outflow rate, episcleral venous pressure and aqueous inflow rate were unaffected by NT, i.e. were the same between NT- and PL-treated cohorts. We then used the modified Goldmann equation (Brubaker 2004) to determine a predicted IOP in the NT-treated eyes and compared this value to the actually measured IOP in these eyes. In brief, Goldmann’s equation states *Q* = *C* (*IOP − EVP*) + *F*_*U*_, where *Q* is aqueous inflow rate, *EVP* is episcleral venous pressure, and *F*_*U*_ is unconventional outflow rate. Under the stated assumptions, the predicted value of IOP in NT-treated eyes is 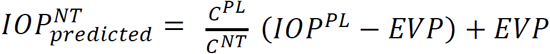, where superscripts indicate whether the value refers to NT- or PL-treated eyes. We assumed a range of EVP values and computed the ratio of the IOP predicted by the above formula to the actual measured IOP value in NT-treated eyes. IOP values used in the calculation were the cohort means of the last measurement taken before sacrifice, as follows: in the prevention study, 17.4 and 25.0 mmHg for NT- and PL-treated eyes, respectively, and in the reversal study, 21.0 and 28.4 mmHg for NT- and PL-treated eyes, respectively.

### Optical Coherence Tomographic Imaging

At the end of the 4 day NT/PL treatment period in the reversal study, and after 4 weeks of NT/PL treatment in the prevention study, OCT imaging was conducted in living mice. In vivo imaging utilized an Envisu R2200 high-resolution spectral domain (SD)-OCT system (Bioptigen Inc., Research Triangle Park, NC). We followed our previously established techniques to image iridocorneal angle structures in mice (Li, Farsiu et al. 2014, Li, Farsiu et al. 2014, Boussommier-Calleja, Li et al. 2015, Meng, Nikolic-Paterson et al. 2016, Li, Lee et al. 2019). Briefly, mice were anesthetized with ketamine (100 mg/kg)/xylazine (10 mg/kg) and maintained with ketamine (60 mg/kg) every 20 min by IP administration. While mice were secured in a custom-made platform, a single pulled glass micro-needle filled with phosphate buffered saline (PBS) was inserted into the anterior chamber of one eye. The micro-needle was connected to both a manometric column to adjust IOP and a pressure transducer (Honeywell Corp., Morristown, NJ, USA) to continuously monitor IOP levels using PowerLab software. The OCT imaging probe was aimed at the nasal or temporal limbus and the image was centered and focused on the SC lumen. While collecting images, mouse eyes were subjected to a series of IOP steps (10, 12, 15, 17 and 20 mmHg) by adjusting the height of the fluid reservoir. At each IOP step, a sequence of repeated OCT B-scans (each with 1000 A-scans spanning 0.5 mm in lateral length) from spatially close positions was captured, registered, and averaged to create a high signal-to-noise-ratio image from the iridocorneal angle region of each animal. The duration of each pressure step was ∼1-2 minutes. Note that the cohort of mice used for OCT imaging was different than that used for facility measurement.

### Segmentation of OCT images

OCT B-scans of iridocorneal angle tissues were registered and segmented following established methods (Li, Mukherjee et al. 2016, Meng, Nikolic-Paterson et al. 2016, Li, Lee et al. 2019) using SchlemmSeg software, which includes two modules: Schlemm I and Schlemm II. Briefly, OCT B-scans were automatically registered using our custom Schlemm I software for SC segmentation. The Schlemm II software package was then used to differentiate SC from scleral vessels, which were automatically marked. If SC was seen to be connected to collector channels (CC), manual separation of SC from CC was required, and was based on the shape of SC and speckling in the images generated by blood cells or other reflectors contained in blood vessels (Mariampillai, Standish et al. 2008, Huang, Zheng et al. 2009, Hendargo, Estrada et al. 2013, Li, Farsiu et al. 2014, Poole, McCormack et al. 2014, Meng, Nikolic-Paterson et al. 2016). The speckle variance OCT-angiography images were generated based on the speckling in SC and vessels as described in detail in previous publication (Meng, Nikolic-Paterson et al. 2016). SC was easily differentiated from other vessels due to its size and location.

### Segmentation Reproducibility

To test the reproducibility of the SC segmentation process, we evaluated both interobserver and intraobserver reproducibility. The segmentation of SC was independently performed by two individuals. The first observer (GL) conducted the experiments and made initial measurements, then repeated the measurements one to two months after the first examination to determine intraobserver reproducibility. The second observer (JC) was first given a training set of images to evaluate, then reviewed the images for the present study in a masked fashion to assess the interobserver reproducibility.

### Inverse FEM Determination of TM Stiffness

In brief, the FEM technique allows one to compute the deformation of a structure due to loads/forces; here we computed the deformation of irideocorneal angle tissues (including the TM) due to changing IOP. In our inverse FEM approach, the stiffness of the TM was parametrically varied until computed SC deformations matched experimental observations, at which point we concluded that the TM stiffness specified in the FEM approach matched the actual (in vivo) value, while masked to treatment group. A pseudo-2D FEM geometry generated in our previous publication (Li, Lee et al. 2019) was used to calculate TM tissue stiffness based on OCT images. We have previously shown (Wang, Johnstone et al. 2017) that such a pseudo-3D approach does not yield results significantly different from a true 3D model, at a fraction of the workload. The model includes the TM, sclera/cornea, and the uvea, and is meshed with 4-noded tetrahedral elements. Tissues are treated as incompressible, isotropic, and nonlinearly hyperelastic (incompressible Mooney*−*Rivlin material model). The TM is assigned a range of stiffnesses (20 kPa to 240 kPa), and for each stiffness value, we simulate the deformation of irideocorneal angle tissues, thus determining the cross-sectional area of SC vs. IOP. The computed SC area is compared with experimental measurements, and the estimated TM stiffness is taken as the value that minimizes the least squares difference between the experimental and predicted normalized SC areas over the IOP range 10 mmHg to 20 mmHg. SC luminal pressure is estimated as previously described (Li, Lee et al. 2019) (Table S1).

### AFM Measurement of TM Stiffness

TM stiffnesses were measured using a previously developed AFM technique on cryosections of mouse eyes (Wang, Read et al. 2017, Wang, Li et al. 2018, Li, Lee et al. 2019). Briefly, de-identified eyes coated with optimal cutting temperature compound (O.C.T.; Tissue-Tek) were cryo-sectioned from 3 different quadrants on a Microm Cryostar NX70 cryostat (Dreieich, Germany). For each quadrant, a few 10 µm thick sagittal cryosections were collected on adhesive slides (Plus gold slide, Electron Microscopy Sciences, Hatfield, PA) and stored for up to 30 min in ice-cold PBS prior to AFM analysis.

Samples were then transferred to an MFP-3D-Bio AFM (Asylum Research, Santa Barbara, CA) and kept continuously immersed in PBS during measurements at room temperature (Figure S5). TM stiffnesses were measured following the same protocol we used previously (Wang, Read et al. 2017). Specifically, cantilever probes were modified by attaching a spherical indenter of diameter 10 µm to smooth nanoscale variations in tissue mechanics. For each indentation, the indentation depth was 0.5–1 µm, with a maximum applied force of 7 nN and approach velocity of 8 µm/s. A Hertzian model was used to extract a Young’s modulus stiffness value from the force versus indentation curves. For each cryosection, the TM was first localized as the region between the pigmented ciliary body and the inner wall endothelium of SC. Multiple locations in the TM region (typically 3–9) were probed by the cantilever and three repeated measurements were conducted at each location. The average from the three measurements was taken as the TM stiffness at that location. TM stiffnesses from all locations within a cryosection were then averaged to obtain the TM stiffness of that cryosection, and values for cryosections within a quadrant were averaged to obtain the TM stiffness in that quadrant. The mean stiffness of all quadrants was taken as the TM stiffness of the eye. Typically, there were 135 force curves acquired per eye (three force curves per location, typically 5 locations per cryosection, and typically 9 cryosections per eye). A small number of measured TM stiffness values were excluded in a post hoc analysis if there was disagreement from a second reviewer as to whether the measurement location lay within the TM.

### Histology, Immunohistochemistry and Transmission Electron Microscopy

After OCT imaging, mice were decapitated under anesthesia, and eyes were collected and immersion fixed in 4% paraformaldehyde at 4°C overnight. The eyes were then bisected, and the posterior segments and lenses were removed. The anterior segments were cut into four quadrants. For immunostaining, each quadrant was embedded into LR-White resin and 1 µm sections were cut and immunostained with antibodies that specifically recognized either alpha-smooth muscle actin (αSMA; 1:100 dilution, rabbit polyclonal, ab5694, Abcam, Cambridge, MA) or fibronectin (FN; 1:50 dilution, mouse monoclonal, Santa Cruz, Dallas, Texas). The secondary antibodies were peroxidase-conjugated AnffiniPure Goat Anti-Rabbit or mouse IgG H&L (Alexa Fluor® 488; Jackson ImmunoResearch Laboratories, West Grove, PA) at 1:500 dilution. Images were captured using a Nikon Eclipse 90i confocal laser-scanning microscope (Melville, NY, USA). Images from NT- and PL-treated eyes were collected at identical intensity and gain settings (Li, Torrejon et al. 2018, Li, Lee et al. 2019). For electron microscopy studies, one quadrant/eye of the anterior segment was fixed in 2% glutaraldehyde and embedded in Epon resin and 65 nm sagittal thin sections were cut through iridocorneal tissues using an ultramicrotome (LEICA EM UC6, Leica Mikrosysteme GmbH, A-1170, Wien, Austria). Sections were stained with uranyl acetate/lead citrate and examined with a JEM-1400 electron microscope (JEOL USA, Peabody, MA).

For quantification of basal lamina deposits below the inner wall of SC, images were captured at 8000× magnification and masked as to the identity of the treatment group. Masked images were scored by two individuals (WDS and CRE) with extensive experience in viewing TEM images of the conventional outflow pathway. Specifically, the density and extent of the extracellular matrix in the basal lamina region was scored on a scale of 0-2, with 0 representing “normal” appearing basal lamina, 2 representing continuous basal laminar deposits such as observed in the mouse steroid glaucoma model (Overby, Bertrand et al. 2014), and 1 representing conditions between the two extremes. Thus, images where included whereby the inner wall of SC was clearly visible, and images were of sufficient quality to examine its basal lamina.

### Statistical analyses

To analyze IOP measurements, the time points from 3-28 days in the prevention study, or the time points from 1-4 days post-NT/PL delivery in the reversal study, were averaged from each eye to produce a single average IOP value per eye for subsequent data analysis. The Mann-Whitney U test was used for analyzing significant difference between groups for IOP and OCT images. To analyze outflow facility measurements, we used the well-established fact that the underlying distribution of outflow facility in mice is log-normally distributed (Sherwood, Reina-Torres et al. 2016). Thus, a weighted paired or unpaired (two-way) t-test (Sherwood, Reina-Torres et al. 2016) was applied to the log-transformed facilities. To analyze data quantifying basal lamina deposits, the Wilcoxon rank sum test was used. Due to that fact that IOPs are normally distributed and the expected directional (lowering only) effect of NT on IOP, we analyzed patient IOP data using a paired t-test with one tail, assuming equal variance. Data in box and whisker plots show median, 25th percentile, and 75th percentile (boxes), as well as minimum and maximum values (whiskers). Data in other plot formats are presented in the form of mean and 95% confidence interval or mean and standard error of the mean (SEM), as noted. A value of *P* ≤ 0.05 was considered statistically significant.

## Acknowledgements

We thank Ying Hao (Duke Eye Center Core Facility), who prepared histology sections and helped with TEM. Dr. Vibhuti Agrahari helped with the preparation and characterization of the NPs. Dr. Sandra Stinnett performed statistical analysis of data quantifying basement membrane deposits and stiffness measurements. We acknowledge funding support from the BrightFocus Foundation, Clarksburg, MA; Research to Prevent Blindness Foundation, NY, NY; the Georgia Research Alliance, Atlanta, GA; Aerie Pharmaceuticals, Durham, NC; and National Institutes of Health, Washington, D.C. (EY030124, EY019696 and EY005722). The sponsors and funding organizations had no role in the design or conduct of this research.

## Author Contributions

Author contributions to the manuscript are as follows: designed research studies (GL, CL, TS, SF, CRE and WDS), conducted experiments (GL, CL, ATR, KW, IR and KY), acquired data (GL, CL, ATR, KMW, RG, IN, JC, KMY, JRS and PC), analyzed data (GL, CL, KW, IN, JC, KMY, RG, TS, SF, JRS, PC, CRE and WDS), provided reagents (ST, SF, CCK, JRS, PC, CRE and WDS), and wrote the manuscript (GL, CL, CCK, CRE and WDS).

## Supplementary Materials

**Figure S1:**
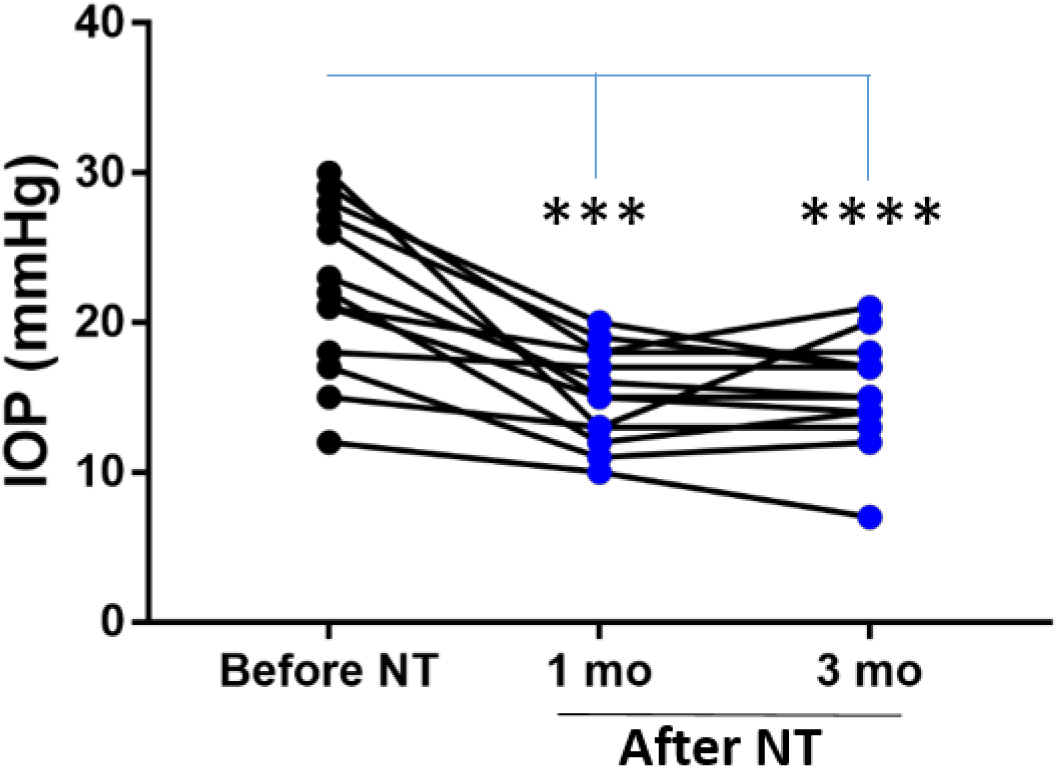
Time course of netarsudil (NT) effects on intraocular pressure (IOP) in patients with steroid-induced elevated IOP poorly controlled with standard glaucoma medications. IOPs were measured by Goldmann applanation tonometry in patients who demonstrated OHT after steroid treatment for a variety of ocular conditions (Table 1). These individuals were initially treated with aqueous humor suppressants and/or prostaglandin analogues but showed persistent OHT, and were thus treated with NT. We show IOPs of the subset of individual patients in cohort 1 that were monitored over 3 months of treatment. “Before NT” indicates IOPs measured before initiating NT in these patients, and “After NT” shows IOP after daily treatment with NT.

**Figure S2.**
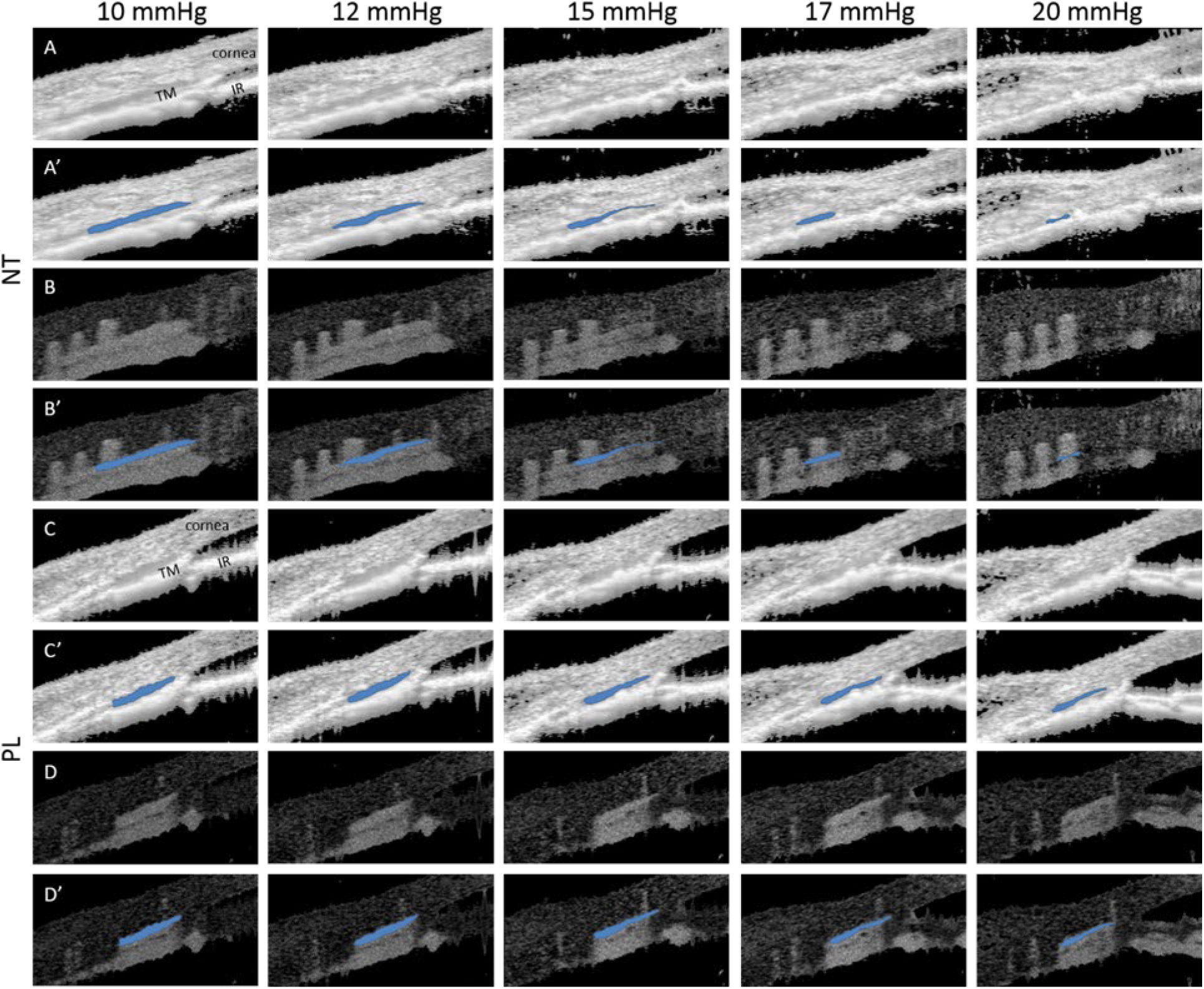
Netarsudil (NT) enhanced IOP-induced collapse of Schlemm’s canal (SC) lumen in ocular hypertensive eyes as visualized by SD-OCT. Mice were treated with DEX-NPs in both eyes for 4 weeks and unilaterally with NT or PL for 4 consecutive days. On day 5, living mouse eyes were cannulated to control IOP and were subjected to sequentially increasing pressure steps to create IOPs of 10, 12, 15, 17, and 20 mmHg. OCT imaging of conventional outflow tissues was conducted in both NT- and placebo (PL)-treated eyes, and the images were averaged to reduce noise at each pressure step. Panel A shows NT-treated eyes, while C shows PL-treated eyes. Corresponding images (A’, C’) show SC lumen segmentation by SchlemmSeg software with lumens indicated in blue. Panels B, B’, D and D’ show the corresponding speckle variance images and SC segmentations. TM, trabecular meshwork; IR, iris.

**Figure S3:**
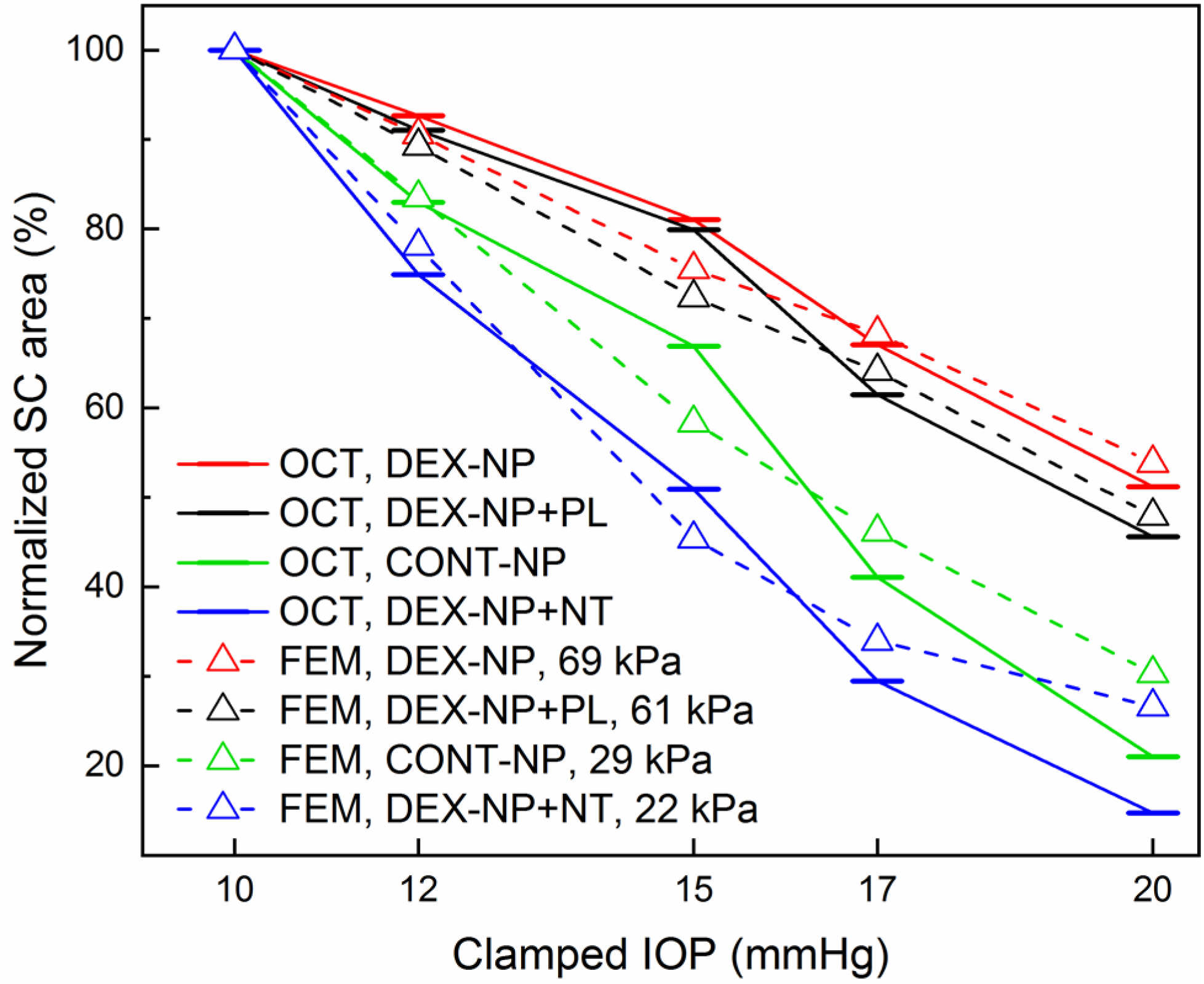
Quantitative comparison of normalized SC lumen areas in NT and PL treatment groups at 5 clamped IOPs (10, 12, 15, 17 and 20 mmHg). This figure is similar to Fig. 3G, but also includes results from eyes with DEX-NP treatment without administration of either NT or PL (DEX-NP, red) and from control eyes that received no glucocorticoids (CONT-NP, green), both adopted from our previous study [44]. The plotted quantity is relative SC area (normalized to value at 10 mmHg) and the bars represent mean values for each IOP. We conducted inverse finite element modeling (iFEM) to structurally analyze the response of anterior segment tissues to varying IOP levels, mimicking the experimental measurements. Dashed lines show results of the inverse FEM analysis for least-squares best fit TM stiffness values, yielding estimated stiffnesses of 61 kPa for PL-treated DEX-NP eyes, 22 kPa for NT-treated DEX-NP eyes, 69 kPa for eyes receiving only DEX-NPs, and 29 kPa for eyes receiving CONT-NPs.

**Figure S4.**
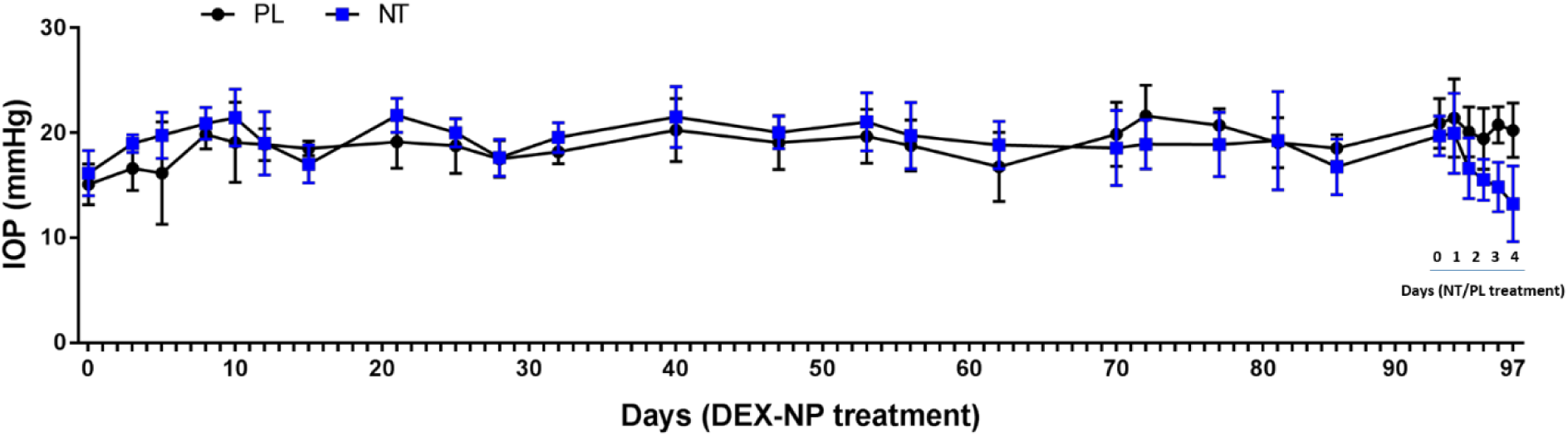
Netarsudil (NT) reversed DEX-NP-induced IOP elevation even after 3 months. Two groups of age- and gender-matched wild-type C57BL/6 mice were treated unilaterally with dexamethasone-loaded nanoparticles (DEX-NPs) twice/month for three months. During the last week, eyes were treated with NT or placebo (PL) once/day for 4 days. IOP was measured at indicated time points. N = 7 animals per group. Notably, this regimen (e.g.: twice/month DEX-NPs) gives a more modest IOP increase (3-4 mmHg) and differs from regimen described for the four week studies in Figure 1.

**Figure S5.**
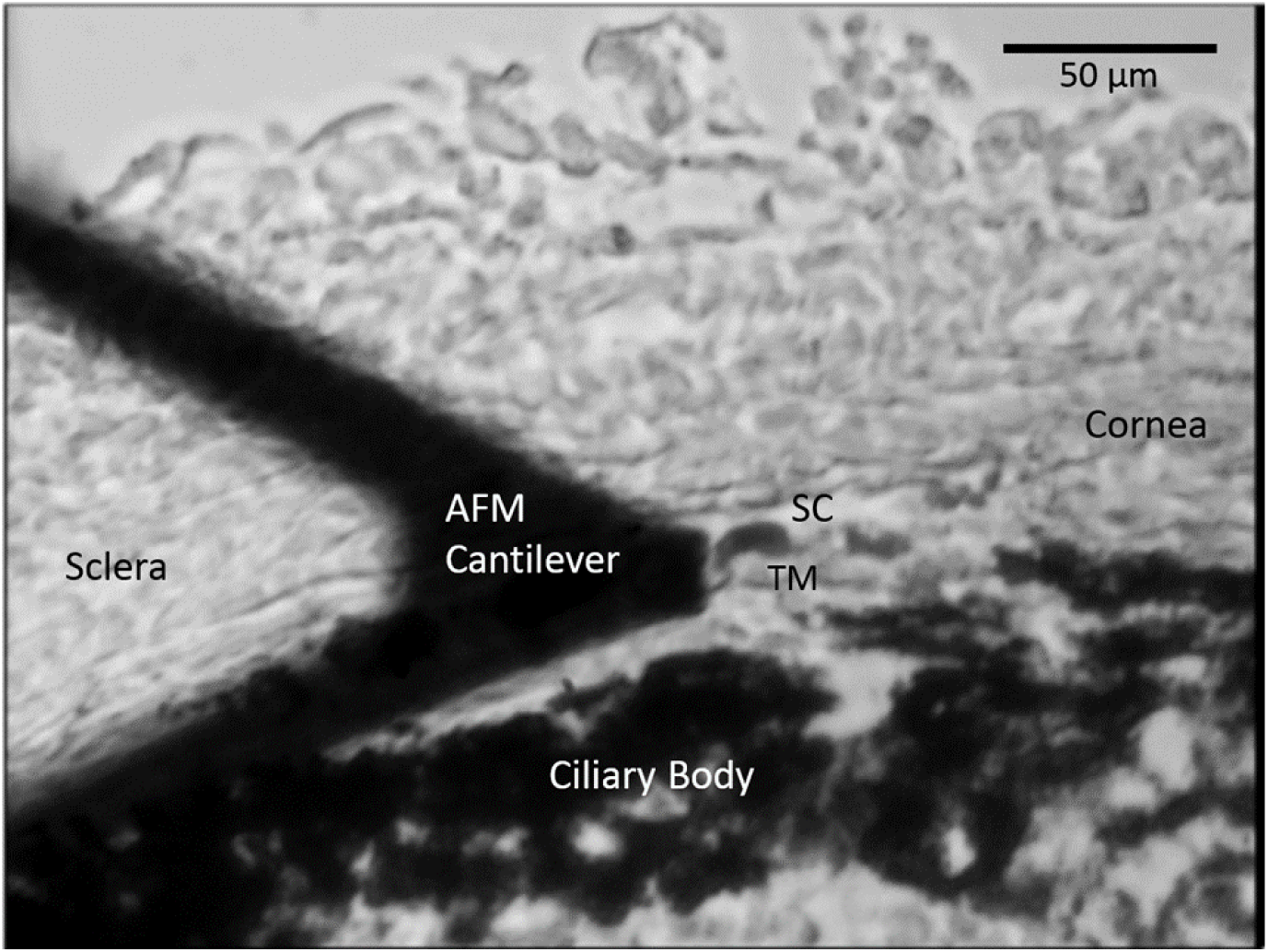
A representative cryosection showing the limbal region during AFM measurements of TM stiffness, acquired on a tissue sample immersed in PBS. AFM, atomic force microscopy, SC, Schlemm’s canal; TM, trabecular meshwork. Section thickness: 10 µm.

**Table S1:** Estimated pressures within SC lumen as a function of clamped IOP levels using two-series resistor model of conventional outflow pathway described previously ^15^.

| Clamped IOP (mmHg) | Schlemm's canal luminal pressure ($P_{SC}$ mmHg) | | | |
| --- | --- | --- | --- | --- |
|  | CON-NP | DEX-NP | DEX-PL | DEX-NT |
| 10 | 7.2 | 6.7 | 6.6 | 7.1 |
| 12 | 8.0 | 7.3 | 7.1 | 7.8 |
| 15 | 9.2 | 8.1 | 7.9 | 9.0 |
| 17 | 10.0 | 8.7 | 8.4 | 9.7 |
| 20 | 11.2 | 9.6 | 9.2 | 10.8 |

IOP: intraocular pressure, PL: placebo, NT: netarsudil, DEX: dexamethasone

